# Taming strong selection with large sample sizes

**DOI:** 10.1101/2021.03.30.437711

**Authors:** Ivan Krukov, Simon Gravel

**Affiliations:** Genome Center, Department of Human Genetics, McGill University, Montreal, Canada

## Abstract

The fate of mutations and the genetic load of populations depend on the relative importance of genetic drift and natural selection. In addition, the accuracy of numerical models of evolution depends on the strength of both selection and drift: strong selection breaks the assumptions of the nearly neutral model, and drift coupled with large sample sizes breaks Kingman’s coalescent model.

Thus, the regime with strong selection and large sample sizes, relevant to the study of pathogenic variation, appears particularly daunting. Surprisingly, we find that the interplay of drift and selection in that regime can be used to define asymptotically closed recursions for the distribution of allele frequencies that are accurate well beyond the strong selection limit.

Selection becomes more analytically tractable when the sample size *n* is larger than twice the population-scaled selection coefficient: *n* ≥ 2*Ns* (4*Ns* in diploids). That is, when the expected number of coalescent events in the sample is larger than the number of selective events. We construct the relevant transition matrices, show how they can be used to accurately compute distributions of allele frequencies, and show that the distribution of deleterious allele frequencies is sensitive to details of the evolutionary model.

## 2 Introduction

The allele frequency spectrum (*AFS*) is an important summary of genetic diversity that is commonly used to infer demographic history and natural selection *(e.g.* Gutenkunst et al. [2009], Kamm et al. [2017], Jouganous et al. [2017]). Given a demographic scenario of population size histories and migrations, these approaches use versions of the diffusion approximation or coalescent simulations to obtain a predicted *AFS.* By comparing predictions to the observed *AFS*’, one can compute likelihoods for different demographic scenarios. These approaches share two drawbacks.

First, the *AFS* calculations can be time consuming for large sample sizes. Some efficient computational shortcuts can be used (e.g., Excoffier et al. [2013], Kamm et al. [2017]) but these often require neutrality. In the presence of natural selection, available methods are more limited, and often involve forward simulations or variants of the diffusion approximation [Gutenkunst et al., 2009, Jouganous et al., 2017].

Second, both Kingman’s coalescent and diffusion-based models rely on an approximation that sample sizes are smaller than the square root of the effective population size 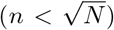, an increasingly problematic assumption [Fu, 2006, Bhaskar et al., 2014]. The assumptions of Kingman’s coalescent can be relaxed to include more general models, involving multiple mergers (e.g., Fu [2006], Spence et al. [2016], and references therein), but accounting for selection in such generalized coalescent models remains challenging.

Because of its ability to handle both complex demographic and selective processes, the diffusion approach has been used to model the distribution of fitness effects for new mutations (*e.g.* Eyre-Walker et al. [2006], Boyko et al. [2008]). However, its validity does not extend to sample sizes used in recent studies that investigate selective constraints [Karczewski et al., 2020].

Jouganous et al. [2017] suggested that a moments-based recursion approach to compute the *AFS* could be modified to include multiple-merger events and selection. Here we derive these recursions and study the impact of large sample sizes on the validity of diffusion models.

We will have to address two challenges. First, recursions are not formally closed under natural selection, meaning that approximations are required. Closure of the moment equations under the neutral Wright-Fisher model occurs because the number of offspring lineages can not be larger than the number of parental lineages. In our notation, the number *n_p_* of distinct parental lineages that are sampled when drawing *n_o_* offspring lineages is at most *n_o_* (see Table 1 for notation throughout). This does not hold under negative selection – selective deaths can require the drawing of *n_p_* > *n_o_* [Donnelly and Kurtz, 1999, Jouganous et al., 2017]. This possible increase in ancestral lineages is also made explicit, for example, in the framework of the ancestral selection graph [Krone and Neuhauser, 1997]. To obtain finite recursions, we will need a closure approximation that can handle strong selection. Second, the additional combinatorial possibilities brought forth by multiple coalescent mergers make the computation of transition matrices for recursion equations more difficult.

**Table 1:** Table of symbols and notation

| Symbol | Meaning |
| --- | --- |
| $N$ | Population size |
| $n_p$ | Number of sampled parental lineages (actual plus rejected samples due to selective deaths) |
| $n_o$ | Number of offspring, sample size |
| $i_p$ | Number of derived alleles in parents |
| $i_o$ | Number of derived alleles in offspring |
| $t$ | Time, in generations |
| $\Phi_{n_o}^t(i_o)$ | Allele frequency spectrum in $n_o$ at generation $t$ |
| $s$ | Selection advantage of the derived allele |
| $r$ | Number of selection events (rejections) in the sample |
| $x_p$ | Frequency of derived allele in parental generation |
| $i_o // n_o$ | A sample with $i_o$ derived alleles “out of” $n_o$ total lineages |
| $\mathbf{H} \begin{bmatrix} i_p // n_p \\ i_o // n_o \end{bmatrix}$ | Hypergeometric probability - sample $i_o // n_o$ in offspring from $i_p // n_p$ in parental generation |
| $\mathbf{T}_r \begin{bmatrix} i_p // n_p \\ i_o // n_o \end{bmatrix}$ | Probability of transition from $i_p // n_p$ in parents to $i_o // n_o$ in offspring, with $r$ selective events<br>- see text for a more detailed description |
| $(i_p, n_p)$ | Event of *drawing* exactly $i_p$ derived out of $n_p$ distinct parental lineages |
| $\mathcal{C}(i_p, n_p)$ | Event of $n_p$ parental lineages *containing* $i_p$ derived alleles |
| $\mathcal{S}_{n_o}(i_o, n_p)$ | Event that $i_o$ out of $n_o$ offspring carry the derived allele and that $n_p$ distinct parental alleles were drawn |
| $\mathcal{S}_{n_o}(i_o, n_p, r)$ | Event that $i_o$ out of $n_o$ offspring carry the derived allele, that the first $r$ draws for the $n_o + 1$th offspring were rejected, and that $n_p$ distinct parental alleles were drawn (across all draws). |

In this article we derive these transition matrices, and show that the additional bookkeeping has the unexpected benefit of making the recursions almost closed.

A mathematical interpretation is that the number of relevant lineages as we go back in time is a random variable that depends on the relative importance of drift (controlled by the population size *N*) and selection (controlled by the selection coefficient *s*), but also on the sample size *n_o_*. The number of in-sample coalescent events per generation scales as the square of the sample size, 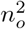, while the number of selective deaths is linear in *n_o_*. Even under strong selection, the expected number of selective deaths is smaller than common ancestry events in sufficiently large samples, meaning that *E*[*n_p_ — n_o_*] ≤ 0. Thus the larger the sample size, the less likely we are to find *n_p_ > n_o_*. Together with an efficient closure approximation for the rare exceptions, this means that strong selection can be handled efficiently and accurately in large samples.

From a practical point of view, we show that these transition matrices can be used to accurately model the distribution of allele frequencies under selection in moderate to large samples, and discuss implications for inference. The implementation of our methods, and the source code for the figures are available at github.com/ivan-krukov/ taming-strong-selection.

## 3 Background

We consider a haploid Wright-Fisher model with *N* individuals per generation, focusing on a single biallelic locus. For a present-day sample with *n_o_* offspring lineages at time *t*, we will be looking for recursion equations for the allele frequency spectrum, 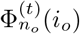, which we define as the probability of observing *i_o_* copies of the derived allele in a sample of size *n_o_*. We will construct this by considering the process of drawing parental alleles (at time *t* — 1) for this finite sample.

To model selection as part of our sampling process, we consider that a selected parental lineage carrying a derived allele can fail to reproduce with probability *s* ≥ 0 (Fig. 1B, broken lines). In such case, we keep sampling until a successful draw. This is equivalent to the usual formulation of the Wright-Fisher model: the probability of drawing a copy of the derived allele is 1 — *s* times the probability of drawing a copy of the ancestral allele. And, because the drawing process depends on the parental allele of both successful and rejected draws, we will see that the *AFS* in the offspring will depend on the state of the *n_p_* distinct parental lineages that were sampled.

**Figure 1:**
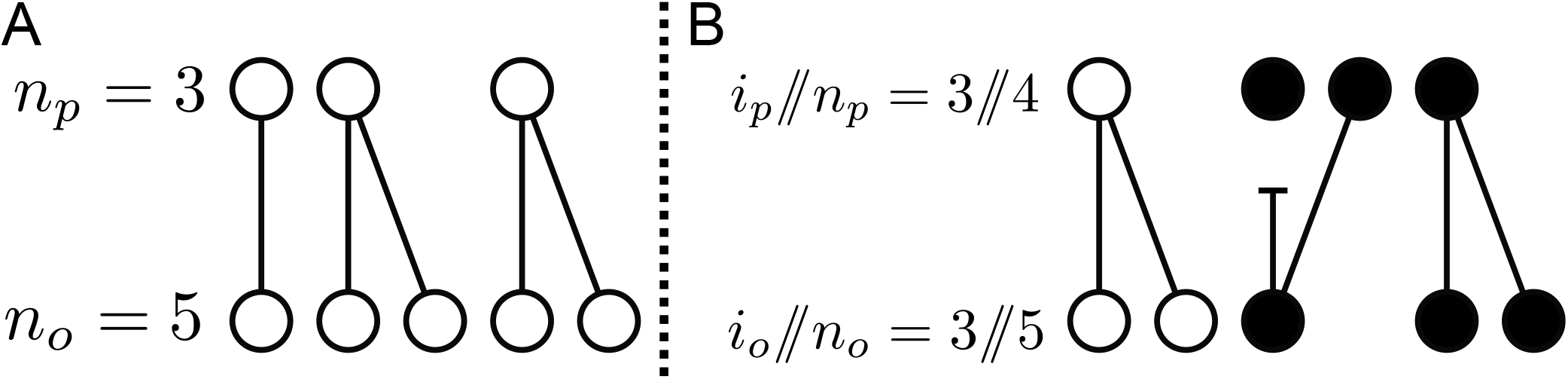
Realizations of sampling parental lineages under neutrality (A) and under selection (B). The top and bottom rows respectively represent the *n_p_* parents and *n_o_* offspring. Filled circles indicate the *i_p_*, or *i_o_* copies of the derived allele; empty circles are ancestral alleles. A line connecting two circles represents a successful draw, a broken line - a selective death triggering a redraw.

To express the allele-frequency spectrum 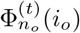 for *n_o_* offspring in terms of the parental *AFS* at *t* — 1, we can sum over possible values for random variables *n_p_* and the number *i_p_* of distinct derived alleles among the parents:

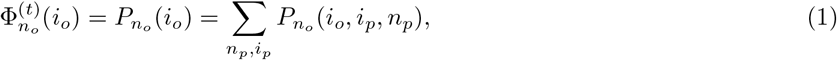

where the subscript *n_o_* indicates dependence on parameter *n_o_*.

To obtain a recursion on the *AFS*, we use an exchangeability property of the parental lineages. Since the order in which we draw new parental lineages in the Wright-Fisher model is random, we can first draw a random permutation of the parental population, and then select unsampled parents in order from this permutation. This allows us to separate properties of this permutation, which depend only on parental *AFS*, from properties of Wright-Fisher sampling, which depend on population size and selection coefficients. Concretely, the event (*i_o_, i_p_, n_p_*) that we draw *i_o_* derived offspring alleles from *n_p_* distinct parents of which *i_p_* are derived can be expressed as the intersection of two events, 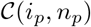 and 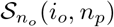. Event 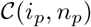 specifies that the first *n_p_* parental lineages from the random permutation carry *i_p_* derived alleles. Event 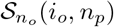 requires that the Wright-Fisher drawing process of *n_o_* offspring drew *i_o_* derived offspring alleles and sampled exactly *n_p_* distinct parental lineages. Event 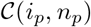 depends on the allele frequency distribution in the parental generation: 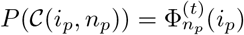 by definition of the *AFS*. Thus

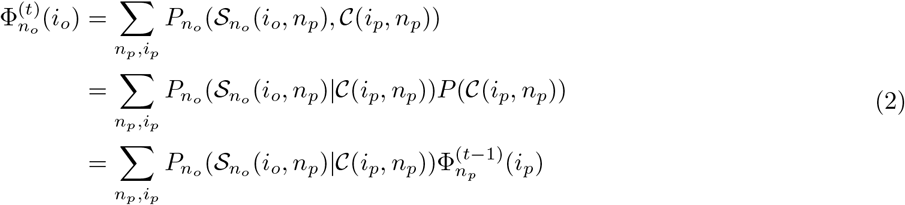

Using matrix notation to account for the sum over *i_p_*, we have

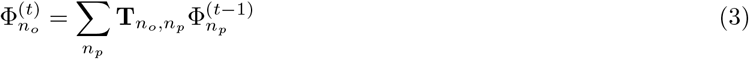

where 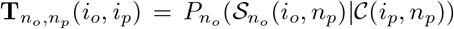 can be thought of as (*n_o_* + 1) × (*n_p_* + 1) transition probability matrix, whose row and column indices correspond to the number of derived alleles in the offspring and contributing parental lineages, respectively. Related recursions under general neutral models were studied in Lessard [2010].

The *AFS* 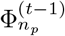, for any *n_p_* ≤ *n_o_* can be obtained by downsampling from 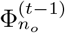 (i.e., 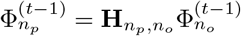 for hypergeometric projection matrix **H***_n_p_,n_o__* if *n_p_* ≤ *n_o_*). Thus, Equation (3) provides a closed form recursion for Φ*_n_o__* under neutrality. This was used in Jouganous et al. [2017] to efficiently compute distributions of allele frequencies under neutrality and in small sample sizes.

Under selection, *n_p_* may be larger than *n_o_*, leading to a set of *N* coupled equations that is typically too large to solve numerically (in the diffusion limit, the number of coupled equations is infinite). But if the number of drift events is typically larger than the number of selective deaths, as happens in large sample sizes, the coupling is weak and we can restore approximate closure by truncating the summation in Eq. 3:

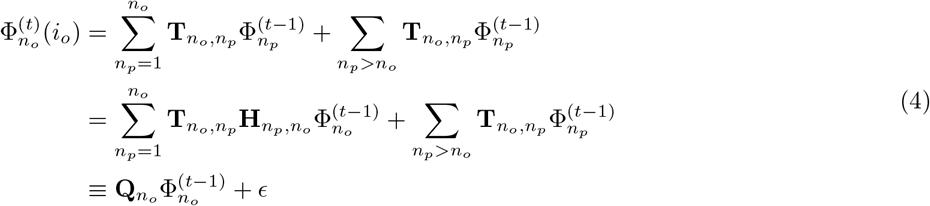

where 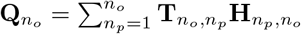 is a square matrix whose (*i_p_, i_o_*)^th^ element is the probability 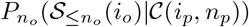 of event 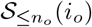 that we observe *i_o_* derived alleles in a sample of size *n_o_* and that we draw *at most n_o_* distinct parental lineages (see Appendix S1.2), given event 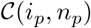 that *i_p_* of the first *n_p_* parental alleles are derived.

The term 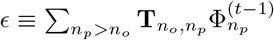 accounts for events with *n_p_ > n_o_*. We will show below that it is small and often negligible for large sample sizes. Further, it can be estimated to ensure convergence even in moderate sample sizes. Using the jackknife extrapolation of [Gravel and NHLBI GO Exome Sequencing Project, 2014], we can write 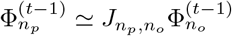 where *J_n_p_,n_o__* is a *n_p_ × n_o_* matrix obtained by jackknife extrapolation, and thus

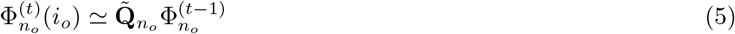

with 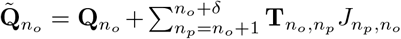 and *δ* is a parameter describing the order of the jackknife approximation to consider.

Jackknife approaches have been used in Jouganous et al. [2017] to derive approximate recursion equations under weak selection. But whereas the approach of Jouganous et al. [2017] requires a jackknife approximation to create a new lineage for every selective death, the formulation in Equation (4) only requires approximations in the (rare) instances where there are more selective deaths than common ancestry events. By accounting for simultaneous events, lineages that go unused due to genetic drift can be repurposed to model selection (Figure 1b). Thus by keeping track of simultaneous events in the transition matrices in (3), we can drastically reduce our reliance on approximations.

Our next goal is to compute these matrices.

## 4 Methods and Results

### 4.1 Constructing the transition matrix

Even though **T***_n_o_,n_p__* in Equation 3 is a combinatorial probability describing a single generation in the Wright-Fisher model, we were unable to compute an analytical expression for it, while simultaneously allowing for multiple coalescences and multiple selective events. However, we can obtain fairly simple recursion by conditioning on the last draw. Such recursions were used for describing large sample size effects without selection in [Bhaskar et al., 2014]. In this section, we provide some mathematical intuition, and a more detailed derivation is provided in the Appendix S1.2.

Under selection, the last draw can be characterized by whether it was successful or rejected, whether it drew a derived or ancestral allele, and whether it drew a previously drawn parental lineage or not (Figure 2). The state of the sampling process prior to the last draw can be described by a number of successfully drawn offspring 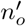, of which 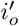 are derived, a number of distinct parental lineages 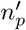 of which 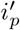 are derived, and a number of failures since last successful draw *r*’. The recursion will be over a probability 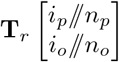 which generalize the transition matrices from Equation (4) to account for rejected draws: 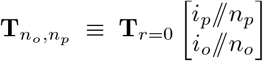. Formally, 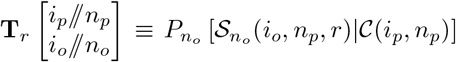 where 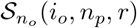 is a generalization of 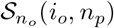 that accounts for rejected draws: it is the event that *i_o_* of the first *n_o_* successfully drawn offspring are derived, that the following *r* draws are rejected, and that the *n_o_* offspring and subsequent *r* failures required exactly *n_p_* distinct parental draws. The bracket notation, 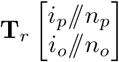, is convenient to keep track of the number of relevant parental lineages (top of bracket) and offspring lineages (bottom of bracket) in the recursion at the cost of obscuring their different probabilistic role.

**Figure 2:**
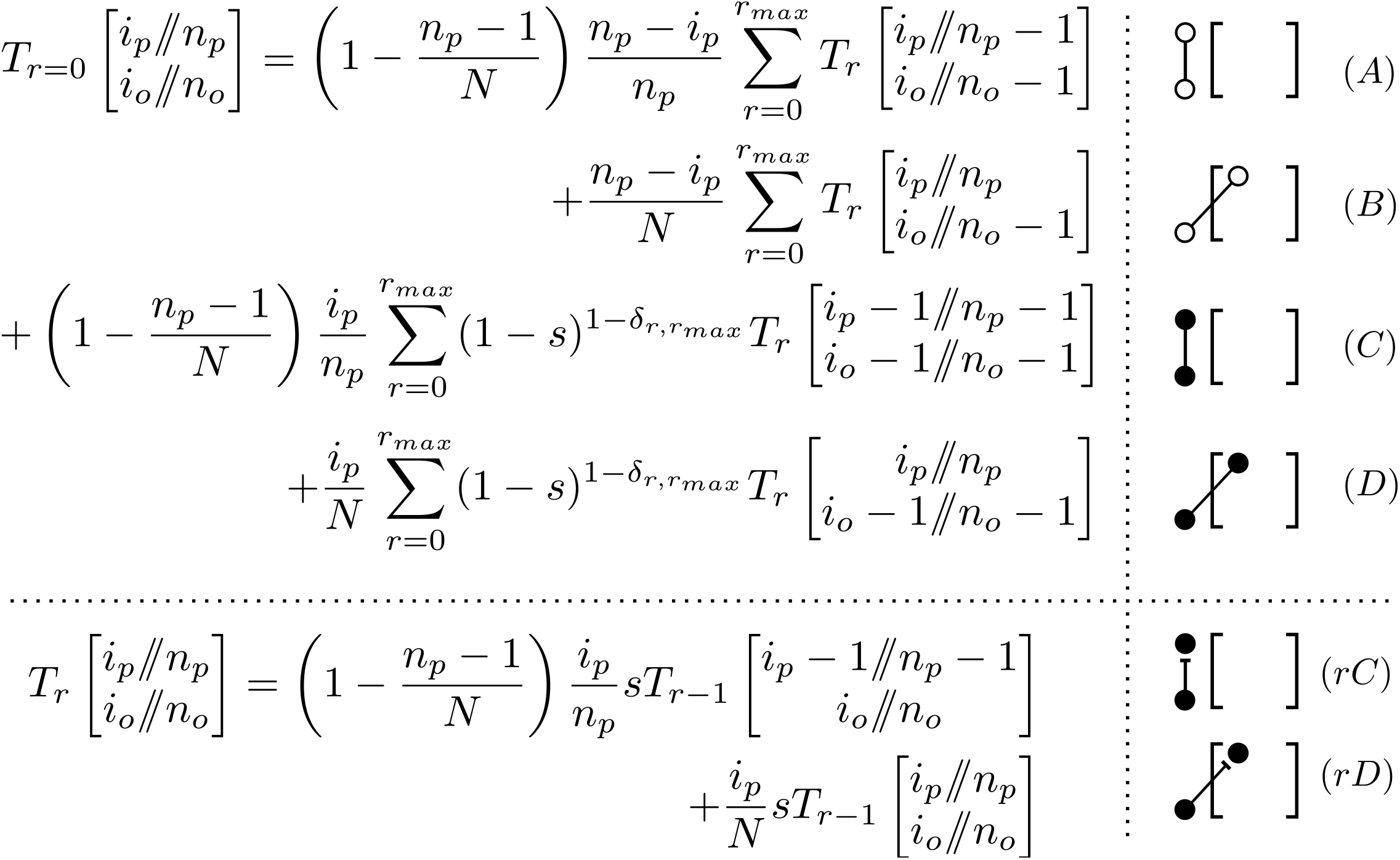
Recursion over the last draw for transition probabilities with selection. The right hand panel represents the last draw graphically. Filled and empty circles represent derived and ancestral alleles. Solid and broken lines are successful lineage draws and re-draws due to selection. Square brackets represent events in sample of size 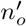 prior to the last draw. Summands *(A-D*) are successful draws where the last lineage is ancestral *(A,B*) or derived (*C,D*). A successful draw can happen after 0 to *r_max_* failed draws, which is accounted for by the sum in the recursion. Terms *rC* and *rD* represent the probability that last draw was rejected. See Table 1 for notation used.

We show in the Appendix and illustrate in Figure 2 that we can express 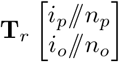 as a sum over a small number of terms of the form 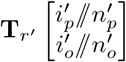. If *r* > 0, the last draw must be a failure and *r*’ = *r* — 1 (leading to cases *rC, rD* in Figure 2). If *r* = 0, the last draw must be a success and 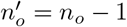 (cases *A* to *D*)

A challenge with this recursion is that the number of rejected lineages *r* leading to each successful draw can be infinite, in principle. However, the probability of having *r* selective deaths for a single successful draw decreases rapidly with *r*, as *s^r^*. We therefore modify the Wright-Fisher model such that at most *r_max_* failed draws *per offspring* are allowed, after which the next draw is immune to selection. Given *s* < 1, we can easily pick *r_max_* to ensure excellent convergence. For example, with *s* = 0.01, which corresponds to *Ns* = 100 with *N* = 10, 000, the probability of having more than three selective deaths is less than 10^-8^, and the probability of having more than eight selective deaths is less than 10^-18^. We have used *r_max_* = 3 in numerical calculations presented below, but a more stringent cutoff does not present particular challenges.

Using caching to avoid re-computing previously computed values, we can systematically compute all the terms in Equation 2. The set of selection transition probability matrices **T** requires 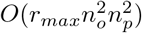 operations to construct: Since each entry in **T**_*r*_ (with *r* > 0) in the recursion requires a constant number of operations, the computational complexity depends on the number of terms we need to calculate. We will need to compute terms of the form 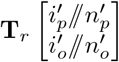, with 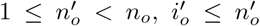, and 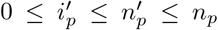, and 0 ≤ *r* ≤ *r_max_*, leading to the bound 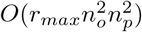.

The terms **T**_*r*=0_ have a bound of the same form (i.e., there are at most 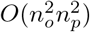 terms, each requiring *r_max_* computations). Finally, the truncated matrix 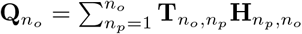 can be constructed in 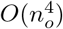.

This is moderately more complex than in the neutral case, where recursions [Bhaskar et al., 2014] and analytical expressions [Lessard, 2010] can be computed in 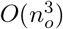.

### 4.2 Calculation of allele frequency spectra

Once the truncated matrix **Q***_n_o__* from Eq. 4 or the jackknife-corrected 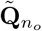 are constructed, they can be used to calculate the allele frequency spectrum under a wide range of models [Jouganous et al., 2017]. To validate the method, we estimate the equilibrium distribution where exact solutions are available [Krukov et al., 2016]. In the infinite sites model at equilibrium, we can compute the equilibrium *AFS,* Φ in a finite sample as a solution to a linear system:

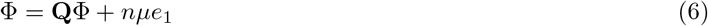

where *μ* is the per-site (forward) mutation rate and *ϵ*_1_ is the second column of the identity matrix of size *n* + 1. Since the infinite-sites model does not account for fixed sites, we only report frequency spectra for polymorphic sites below.

Figure 3 shows the comparison of the *AFS’* calculated from Equation 6 with a jackknife of order 5, the diffusion approximation [Ewens, 2004, eq. 9.23], and the numerical calculation performed in `moments` [Jouganous et al., 2017]. In all the panels, the sample size is *n* = 200 individuals. Any allele frequency probabilities below 1 × 10^-12^ are not included in the figures.

**Figure 3:**
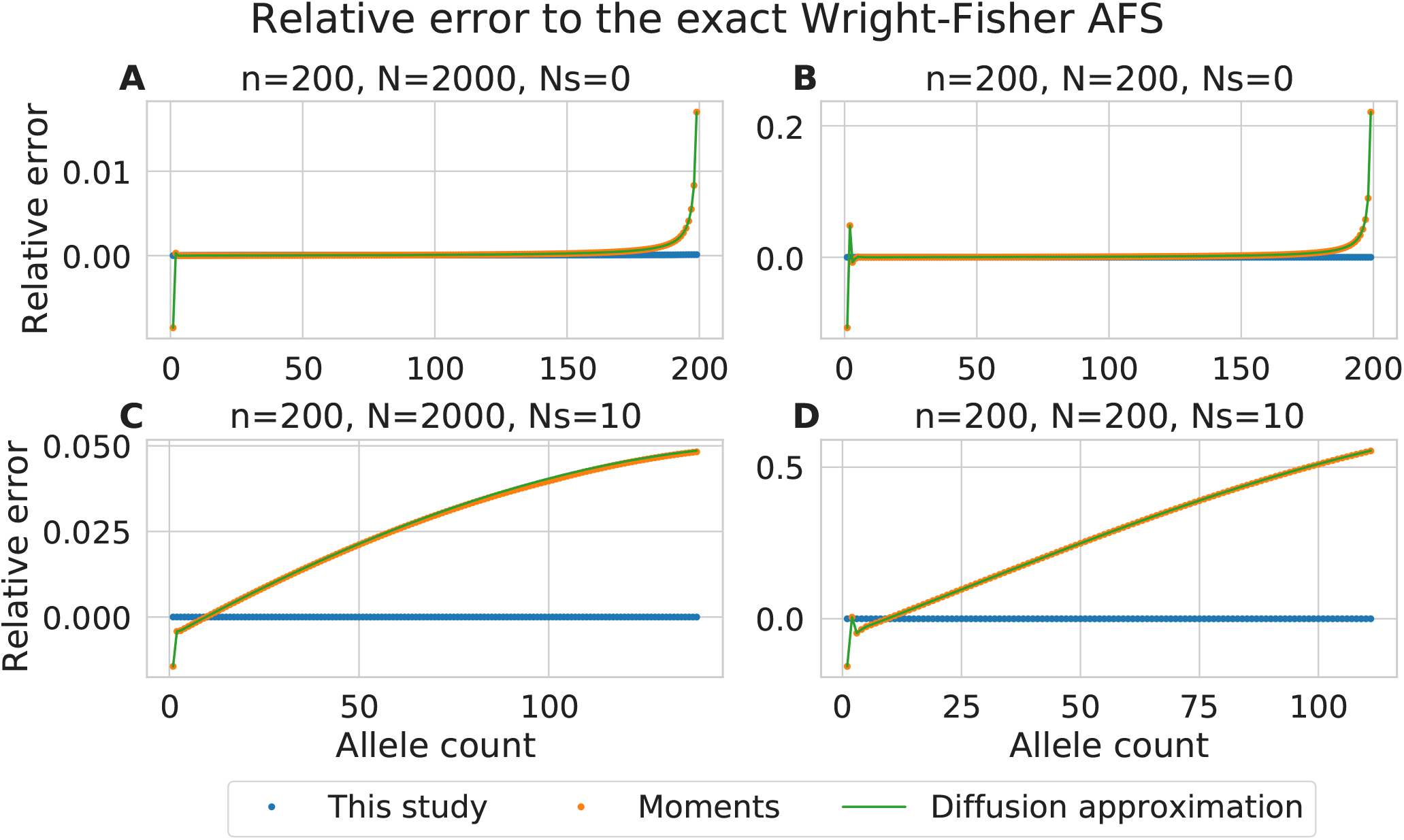
Relative error of allele frequency spectra in a sample of size *n* = 200 with respect to the Wright-Fisher model. Neutral (*Ns*=0) (A,B), or deleterious alleles (*Ns* = 10) (C,D). (A) shows the frequency spectrum in a sample from a large population (*N* = 2000), (B) The sample is entire population (*n = N* = 200). At neutrality all the models agree, except at high allele frequencies. With negative selection, the predictions differ substantially at moderate allele frequencies. Note that *Y*-axes are at different scales between the panels. An extended version of the figure is shown in the appendix - figure S2.

*AFS* computed using the recursion with the jackknife (Figure 3) are indistinguishable from the exact Wright-Fisher solution *AFS* in all the examined parameter ranges. *AFS* computed using the present approach without the jackknife (i.e., with pure truncation, *ϵ* = 0, Figure S1) show moderate departures from the exact results, mostly at high allele frequencies where the number of observations is low. Distributions of allele frequencies using `moments` with a jackknife shows excellent agreement to predictions of the diffusion approximation for small selection coefficients, but visible discrepancies with *Ns* ≥ 50. Thus selection is treated more accurately in the present approach, as expected.

However, the larger difference between the two approaches stems not from is the difference in the accuracy of the jackknife approximations but in the contrast between diffusion-based and Wright-Fisher-based genealogies.

Figures 3A,B show that there is a disagreement between the exact Wright-Fisher and the diffusion solutions for singletons, doubletons, and (*n* — 1)-tons even under neutrality. This effect increases with sample size, and has been observed by comparing Kingman’s coalescent with multiple-merger coalescent events in [Fu, 2006, Bhaskar et al., 2014].

Figures 3C,D show a comparison at moderate negative selection, *Ns* = 10. Here, we see a stronger difference between diffusion and Wright-Fisher models. Comparisons under more parameter choices can be found in Figure S2.

Because differences between diffusion and Wright-Fisher models are tied to multiple merger events, we do not expect large differences in intensive quantities such as the genetic load, and this is confirmed in Supplementary Table S1. Rather, the distribution of allele frequencies in Figure 3 indicates that a higher proportion of the load is caused by common variants under the diffusion model relative to the Wright-Fisher model.

This excess of high-frequency deleterious variants in diffusion models suggests a difference in the rates of fixation of deleterious alleles. Figures S3 and S4 show that these differences remain modest except for fairly deleterious alleles and small populations, where the differences are large enough to affect mean fitness over evolutionary time-scales.

When the sample is the entire population, as in figures 3 B and D, the present method is exact. It is also not needed, since exact recursion equations can be obtained from binomial sampling *(i.e.,* there is no computational benefit to consider a subsample that is as large as the entire population). The present approach is useful where accurate recursions can be obtained for sample sizes that are appreciably smaller than N. We therefore seek asymptotic results for the convergence of the recursion approach in the following sections.

### 4.3 Estimating the missing probability

Given that 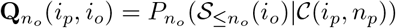, we can compute the fraction of unaccounted-for or approximated draws due to truncation: Given *i_p_* derived among the first *n_o_* parental alleles, this is 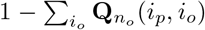. From a numerical perspective, this allows for easy tracking of the proportion of the probability transition that required a jackknife approximation. Figure S5 shows the probability of missing lineages under the worst-case scenario where all parental alleles are derived, *i_p_* = *n_p_*. The probability first increases with increasing sample size, as the number of draws that can result in selective deaths increases linearly. However, the number of drift events eventually overtakes the number of selective events, and the probability that we need additional lineages decreases rapidly with sample size. Creating additional lineages via the jackknife approximation improves the performance further.

Supplementary section S1.4 provides analytical expressions for the probability distribution *P*(*n_p_ — n_o_*) under worst-case scenario *i_p_ = n_p_*, as well as its mean and variance. In particular, we show that

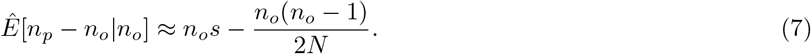

We thus have the usual interplay of selection and drift. Solving for 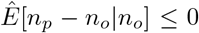 yields a critical sample size 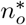 where there are more drift events than selection events in the sample,

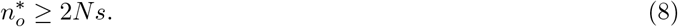

We also show that, to leading order in *s, n_o_/N*, and 1/*N*, the variance is

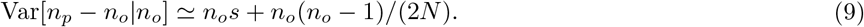

Thus the mean and variance to leading order are those of the Skellam distribution: the *increase* in lineages can be modelled, to lowest order, as the number of lineages gained through selection (modelled as Poisson with mean *n_o_s*) minus the number of lineages lost to drift (modelled as an independent Poisson variable with mean *n_o_*(*n_o_* — 1)/(2*N*)). As the number of drift and selection events get larger, the Skellam distribution can be approximated as a Gaussian with corresponding mean and variance.

Figure S9A shows the excellent agreement between the exact distribution and the Gaussian approximation (using the exact variance from Equation 27). By contrast, we don’t expect this approximation to work well if the sample size is so small that few drift events occur at a given generation, (say, 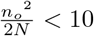), where the Skellam distribution is a better fit - Figure S7.

The Gaussian approximation using leading order terms provides intuition about regimes where we expect drift to almost always overtake selection. Appendix S1.6 shows that the sample size *n_z_* required such that the expected loss in the number of lineages is *z* times its standard deviation obeys

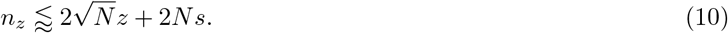

The second term encodes the condition that there are more coalescence events than selection events, on average. It is comparable to contemporary sample sizes even under strong selection (say, 2*Ns* = 100) The first term is independent of s and can be interpreted as a condition that there are sufficient drift events per generation to ensure that the probability of having zero drift event is weak: 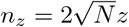 implies 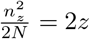. One interpretation is that we need at least a few expected coalescence events per generation to ensure that every selective event is compensated by a coalescence event at the same generation.

For a fixed *z* = 3, Figure S9B shows the critical sample size relative to the population size (*n_c_/N*). Together with equation 10, this shows that the truncation can be accurate even for strong selection and sample sizes much smaller than the full population size.

If we extrapolate the Poisson model of the drift vs selection interplay in equations (7) and (9), we can approximate the critical sample size for a given parental frequency *x_p_* as

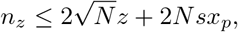

suggesting that even very strong selection will not overcome drift even in moderate sample sizes.

Thus a determinant of the accuracy of the present approach is not whether selection is weak enough, but whether there is enough drift in our sample to ensure that each selection event is met by a drift event in the same generation.

### 4.4 Integrating over several generations

This naturally suggests an improvement: if we consider transitions in the *AFS* not from one generation to the next, but from one generation to *g* generations down the line, drift at one generation can compensate for selection events at another. If the *n_o_* offspring depend on *n_p_* = *n_o_* + 1 distinct parental lineages, because a selective event occurred, but these parental alleles depend on only *n_gp_* = *n_o_* grand-parental lineages, because of a drift event, a transition matrix from the grand-parental AFS to the offspring AFS would be closed.

Such a transition matrix require additional bookkeeping, summing over possible intermediate parental configurations. As it turns out, computing accurate transition matrices over a few generations is reasonably straightforward and not significantly more costly than computing single-generation matrices (Appendix S1.7). If the expectation of 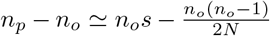 is negative, the probability that we require more than *n_o_* ancestral lineages *g* generations ago decreases with *g*. To get an idea of the truncation behavior for small number *g* ≪ *N* of generations, we can approximate the change in the number of lineages as we go back in time as a random walk with (approximately) constant mean and variance. In this case, both the mean and variance will be scaled by a factor *g* relative to the single-generation case, and Equation (10) becomes:

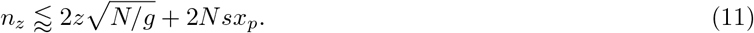

For example, consider a population of size *N* = 10, 000 with strong selection of 2*Ns* = 100. If we set *z* = 5 and *g* = 10, for example, to reach high accuracy even without a jackknife, we require a sample size of at most 400 = 0.04*N*.

As long as *n* > 2*Nsx_p_*, we can choose a number of generations to ensure that the probability of *n_p_ > n_o_* is small.

## 5 Discussion

The diffusion approximation and Kingman’s coalescent models differ from the Wright-Fisher model in that they do not account for possible simultaneous events. A common interpretation is that they are approximations to the ‘exact’ Wright-Fisher process (e.g., Fu [2006]). In this interpretation, the approach presented here provides an overall more accurate way of estimating distributions of allele frequencies for deleterious variants in large samples than the state of the art.

Under neutrality, where differences across models have been studied extensively, the coalescent remains useful as an approximation even in rather large sample sizes [Fu, 2006], although quantitative inference can be affected in realistically large samples [Bhaskar et al., 2014]. Overall, we find that the same holds under selection in large populations and modest selection coefficients, with differences being exacerbated and reaching qualitative levels in small populations and large selection coefficients (Figures 3 and S2).

Whereas differences among neutral models were restricted to the extremes of the distribution, differences under selection span the entire distribution of allele frequencies (Figure 3). Such discrepancies point to differences in the distribution of gene genealogies that go beyond details of whether coalescence events are simultaneous or sequential within a generation. Fu [2006] proposed a rather detailed comparison of gene genealogies under both (neutral) models.

To directly compare how within-generation differences affect long-time gene genealogies, we consider a discretized version of Kingman’s coalescent in which the times of all coalescence events are rounded up to the previous generation. This discretization turns Kingman’s coalescent into a Cannings model that has simultaneous and multiple mergers, but whose gene genealogies have, up to rounding, the same distribution as in Kingman’s coalescent.

Figure 4 shows a comparison of sibship sizes under discrete Kingman coalescent and Wright-Fisher model, in a sample equal to the population of size *n* = *N* = 100000. The discretized coalescent has slightly larger families overall, (i.e., the rate of coalescence is slightly higher, as has been observed e.g. in Fu [2006]), but the differences in sibship sizes are not uniform: There are a few more sibships of size 1 and many more families of size above 4 in a coalescent model, whereas there are more parents with 0, 2, or 3 offspring under Wright-Fisher. Under neutrality, the excess of single-child families with the coalescent leads to an excess of deep singleton lineages and thus (*n* — 1)-tons, as discussed in Fu [2006]. The abundance of large families within the coalescent provides a higher probability for new mutations, including deleterious ones, to rapidly reach appreciable frequency in the populations.

**Figure 4:**
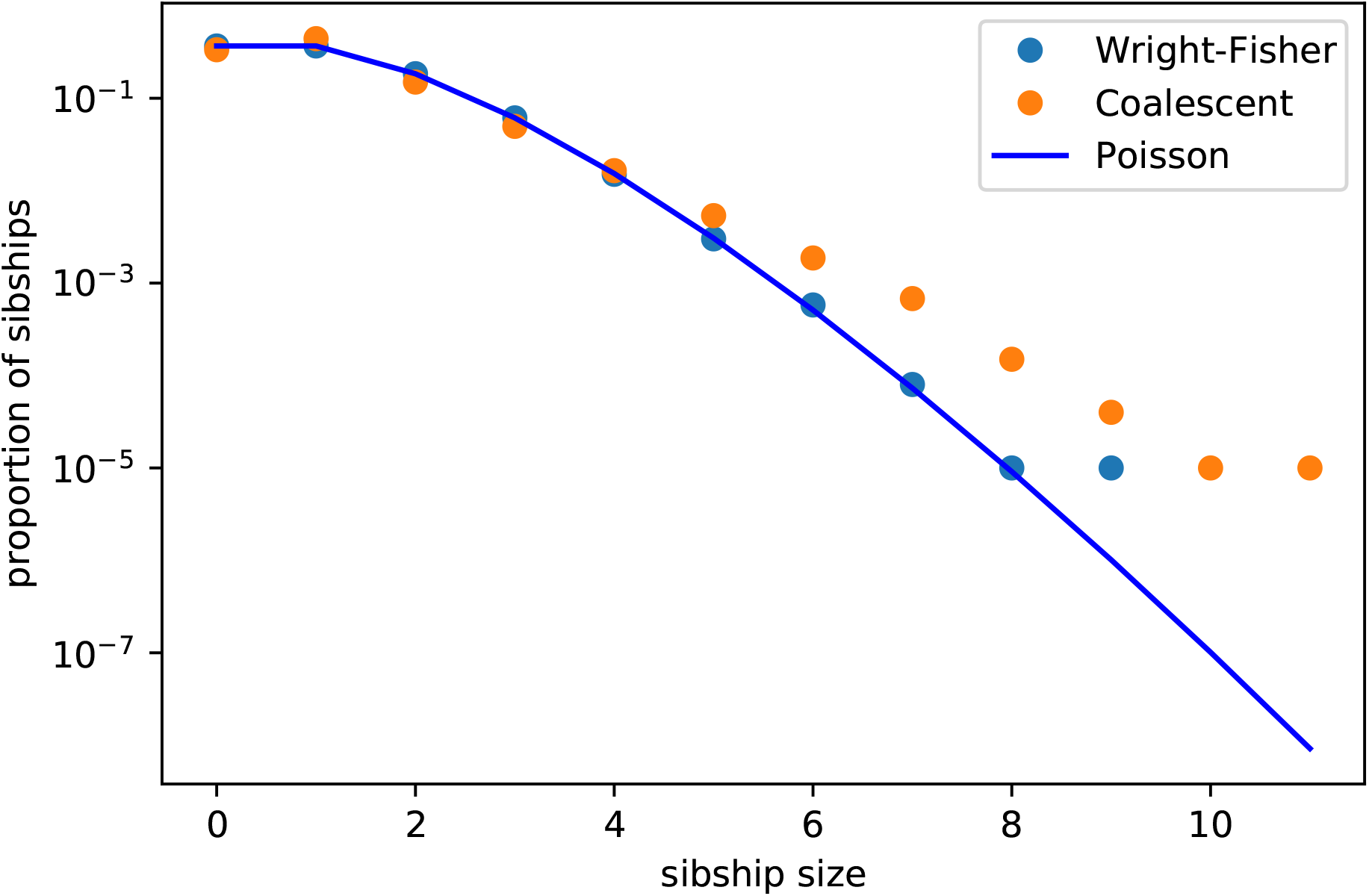
Distribution of family sizes under Kingman’s coalescent and Wright-Fisher model, as simulated under `msprime`. Nelson et al. [2020] in a population of size 100,000. In Kingman’s coalescent, the sibships with zero siblings are obtained by computing *N* — *n_p_*. The solid line shows the Poisson distribution.

The larger differences in prediction of allele frequency distributions between Wright-Fisher and Kingman’s coalescent models under selection is a bit of a concern for inference, as neither approach provides a particularly accurate model of the distribution of family sizes in real populations. If our inferences depend strongly on the differences between family sizes under the two models, we can expect that both models will perform poorly.

As much as we would like to, we cannot state with confidence that recursions under the Wright-Fisher model presented here are a more faithful representation of natural gene genealogies than those based on the coalescent. However, they do present substantial advantages.

First, the handling of natural selection is more robust than in existing moments-based approach, as most selective events are now treated exactly rather than using an empirically accurate but uncontrolled jackknife approximation.

Second, the approach can be generalized to more robust models so that we are no longer bound to the diffusion approximation. Lessard [2010] has derived recursions for a wide range of neutral models, and we expect that many of those, including the ancestral selection graph, can be generalized to almost-closed recursions with selection.

Finally, even though we focused here on simple demographies, many methods for the analysis of complex demographies, such as `momi` [Kamm et al., 2017], build on discrete transition matrices for the *AFS* but have been limited to neutrality. The existence of robust transition matrices under selection opens up the possibility of extending these methods to handle natural selection.

A remaining challenge of the present approach is that the computation of the transition matrices is itself costly. It can be pre-computed, but this requires step-wise constant population sizes and selection coefficients. Efficient implementation of this approach for continuously-varying population sizes and selection coefficients, in the style of the `moments` software, will require some additional analytical work.

One of the main benefits of studying large sample sizes of whole-genome genetic data is to refine our understanding of strongly deleterious variation [Karczewski et al., 2020]. Performing quantitative inference for such datasets will require models that can handle both strong selection and large sample sizes. Since the transition matrices defined here can be used in both forward [Jouganous et al., 2017] and backward approaches [Kamm et al., 2017], they can help genetic models catch up with the requirements of genetic data.

## 6 Acknowledgements

We thank Sabin Lessard and Aaron Ragsdale for useful discussions.

## S1 Appendix

### S1.1 Additional figures

#### S1.2 Deriving the recursion on the transition matrices

The transition matrices 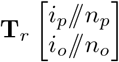 are defined in terms of sampling probabilities 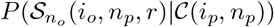, where 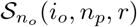 is the event that *i_o_* of the first *n_o_* offspring carry the derived allele, that the r draws following the *n_o_*th drawn offspring are rejected, and that at the end of the *r*th rejection we have drawn exactly *n_p_* parents. Our goal here is to derive a recursion over the last event *ℓ*. We characterize the last draw event in terms of whether it is successful (*σ_ℓ_* = 1) or not *(σ_ℓ_* = 0); whether it draws a derived allele *(γ_ℓ_* = 1) or not *(γ_ℓ_* = 0), and whether it draws a previously undrawn parental allele (*δ_ℓ_* = 1) or not (*δ_ℓ_* = 0).

**Table S1:** Genetic load under the allele frequency spectra from present study (*L*), and the diffusion approximation (*L_diffusion_*), with sample size *n* = 200 individuals. Load calculated as 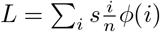, relative error is 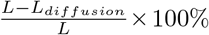.

| N | Ns | $L$ | $L_{diffusion}$ | Relative error |
| --- | --- | --- | --- | --- |
| 200 | 0 | 0 | 0 | 0% |
| | 1 | $1.149e-8$ | $1.150e-8$ | 0.086% |
| | 5 | $1.128e-8$ | $1.130e-8$ | 0.181% |
| | 10 | $1.054e-8$ | $1.056e-8$ | 0.148% |
| | 20 | $1.025e-8$ | $1.026e-8$ | 0.140% |
| | 50 | $1.009e-8$ | $1.010e-8$ | 0.146% |
| 400 | 0 | 0 | 0 | 0% |
| | 1 | $1.150e-8$ | $1.150e-8$ | 0.041% |
| | 5 | $1.129e-8$ | $1.130e-8$ | 0.090% |
| | 10 | $1.055e-8$ | $1.056e-8$ | 0.073% |
| | 20 | $1.026e-8$ | $1.026e-8$ | 0.068% |
| | 50 | $1.010e-8$ | $1.010e-8$ | 0.068% |
| 1000 | 0 | 0 | 0 | 0% |
| | 1 | $1.150e-8$ | $1.150e-8$ | 0.016% |
| | 5 | $1.130e-8$ | $1.130e-8$ | 0.036% |
| | 10 | $1.056e-8$ | $1.056e-8$ | 0.028% |
| | 20 | $1.026e-8$ | $1.026e-8$ | 0.027% |
| | 50 | $1.010e-8$ | $1.010e-8$ | 0.026% |
| 2000 | 0 | 0 | 0 | 0% |
| | 1 | $1.150e-8$ | $1.150e-8$ | 0.009% |
| | 5 | $1.130e-8$ | $1.130e-8$ | 0.017% |
| | 10 | $1.056e-8$ | $1.056e-8$ | 0.014% |
| | 20 | $1.026e-8$ | $1.026e-8$ | 0.014% |
| | 50 | $1.010e-8$ | $1.010e-8$ | 0.013% |

**Figure S1:**
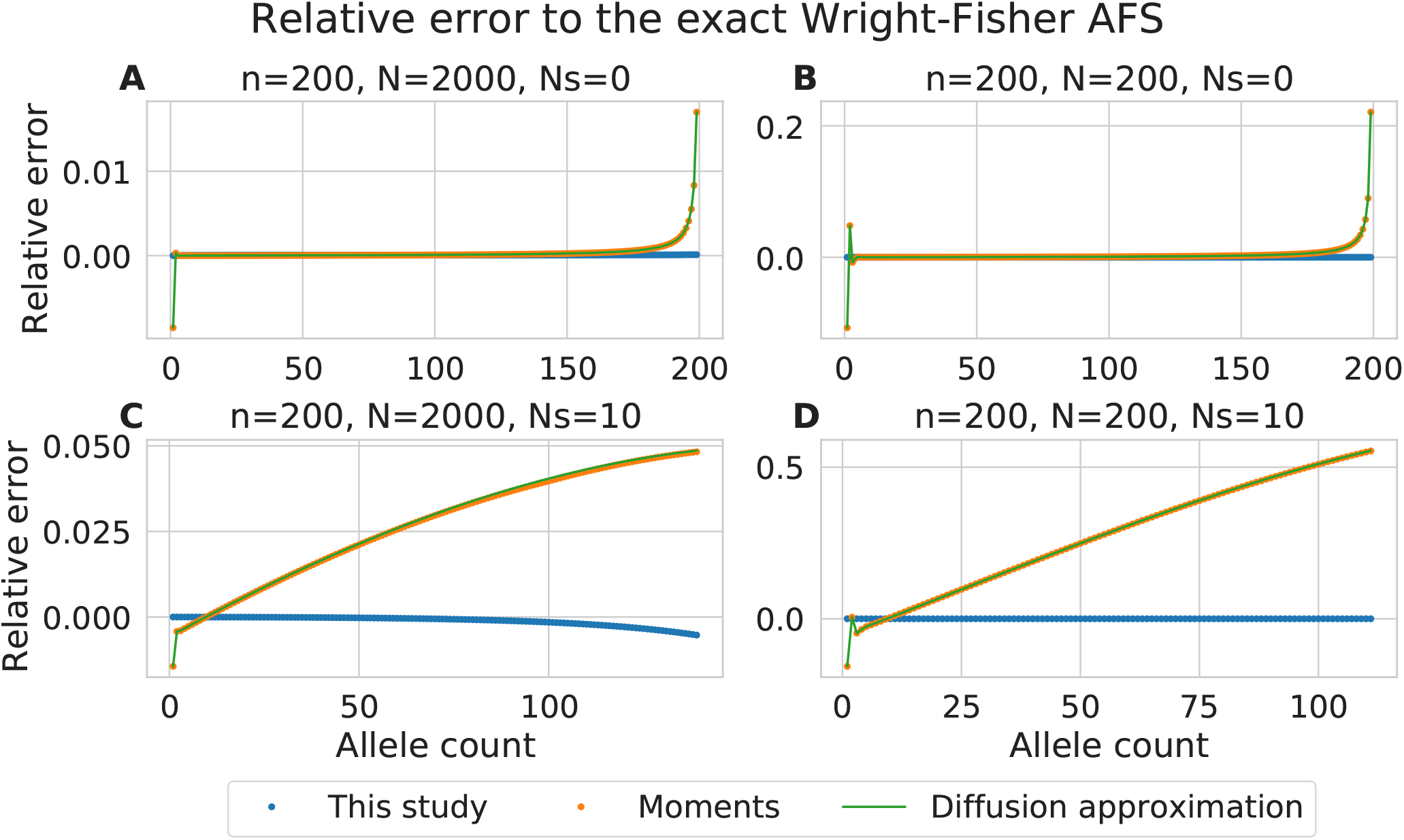
Allele frequency spectra calculated in Figure 3 without the use of the jackknife.

With this notation in place, we can write our recursion as:

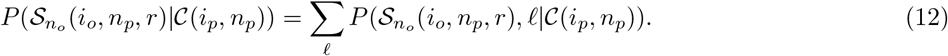

Now that we have explicitly specified the last draw *ℓ*, we can remove the corresponding information from event 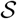,

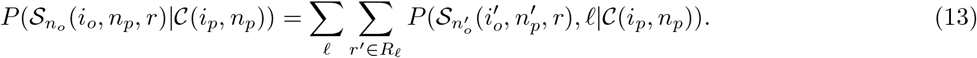

where 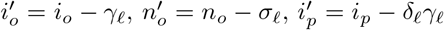, and 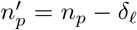 represent the counts of derived and total alleles among the offspring and parents prior to the last draw. The number of failures *r*’ can take more than one value if the last draw was a success, so that

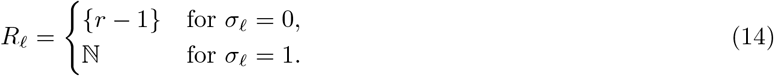

We can further simplify the notation by defining shorthand for events prior to the last draw, 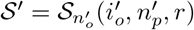, and 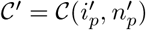, and after the last draw 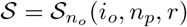, and 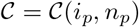

The event 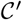 is fully determined by 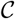 and *ℓ*, so that we can write

**Figure S2:**
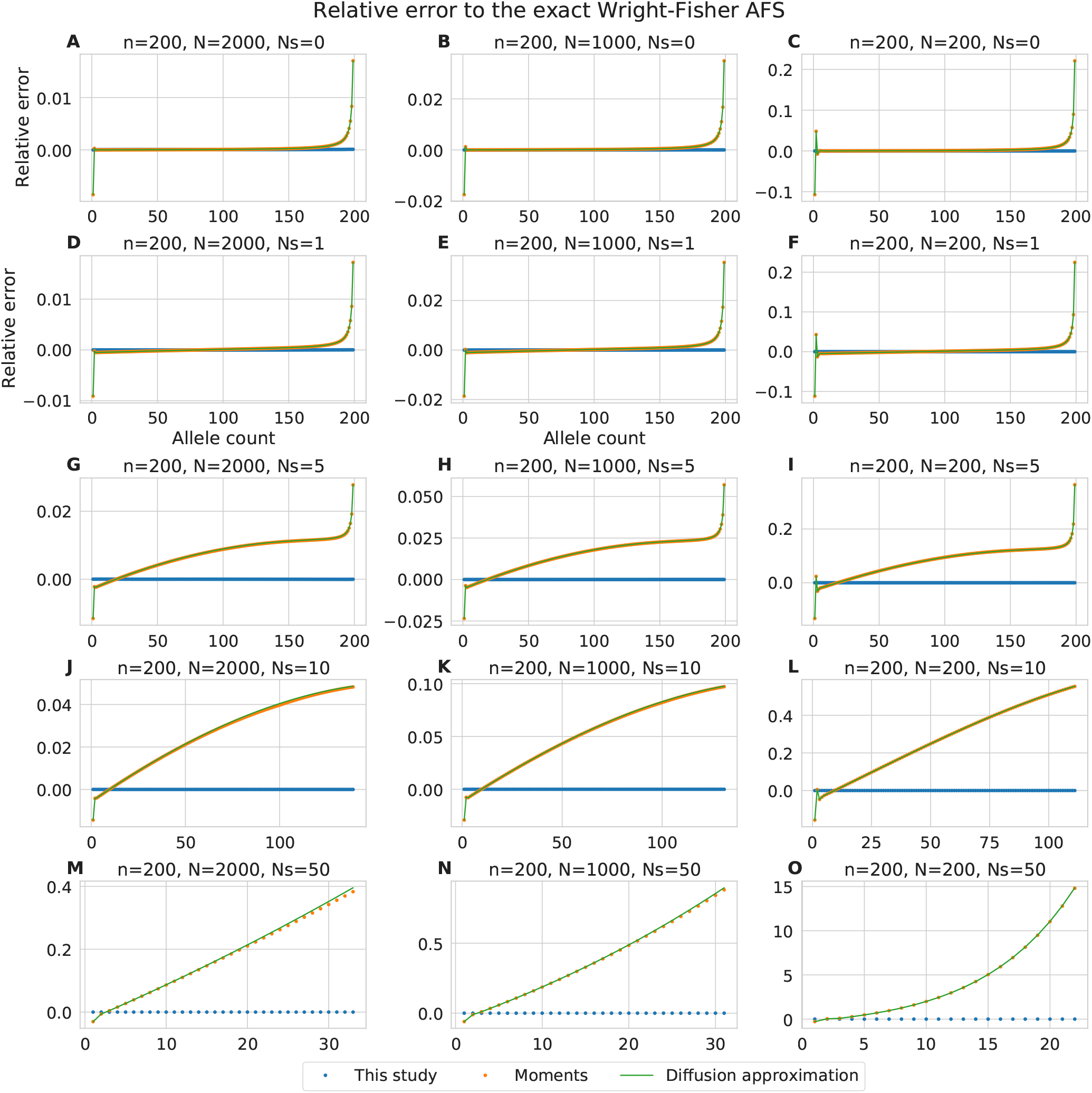
Extended version of figure 3. Relative error of allele frequency spectra from different models with respect to the Wright-Fisher *AFS.* Panels horizontally - population size (*N* = 2000, 1000, 200). Note that the last column corresponds to the special case where the sample is an entire population. Panels vertically - selection coefficients (*Ns* = 0,1, 5,10, 50). Note that in the strong selection case (*Ns* = 50), the *X*-axis is truncated, such that the total probability of an allele at any frequency is above 1*e* — 12.

**Figure S3:**
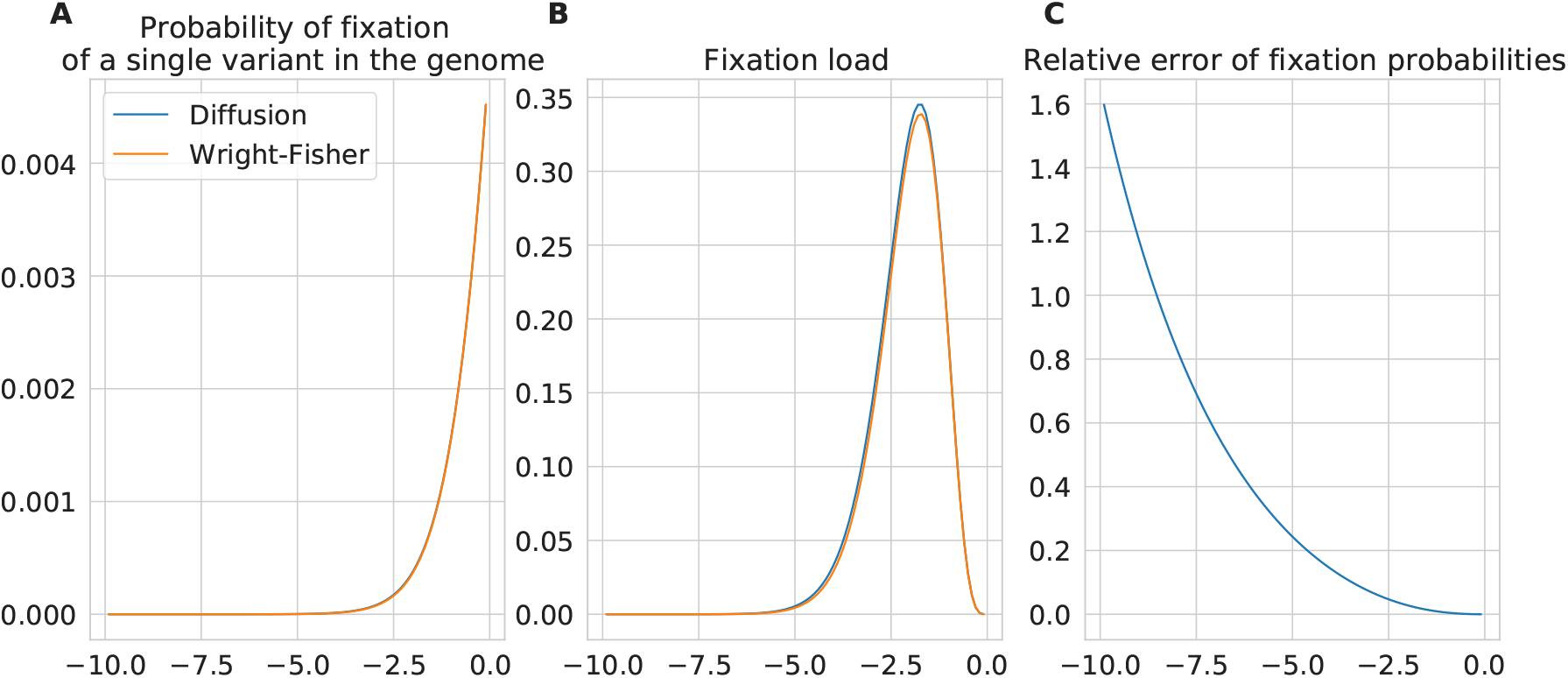
(A) Fixation probability, (B) fixation load, and (C) relative error between the diffusion and Wright-Fisher models for N=100.

**Figure S4:**
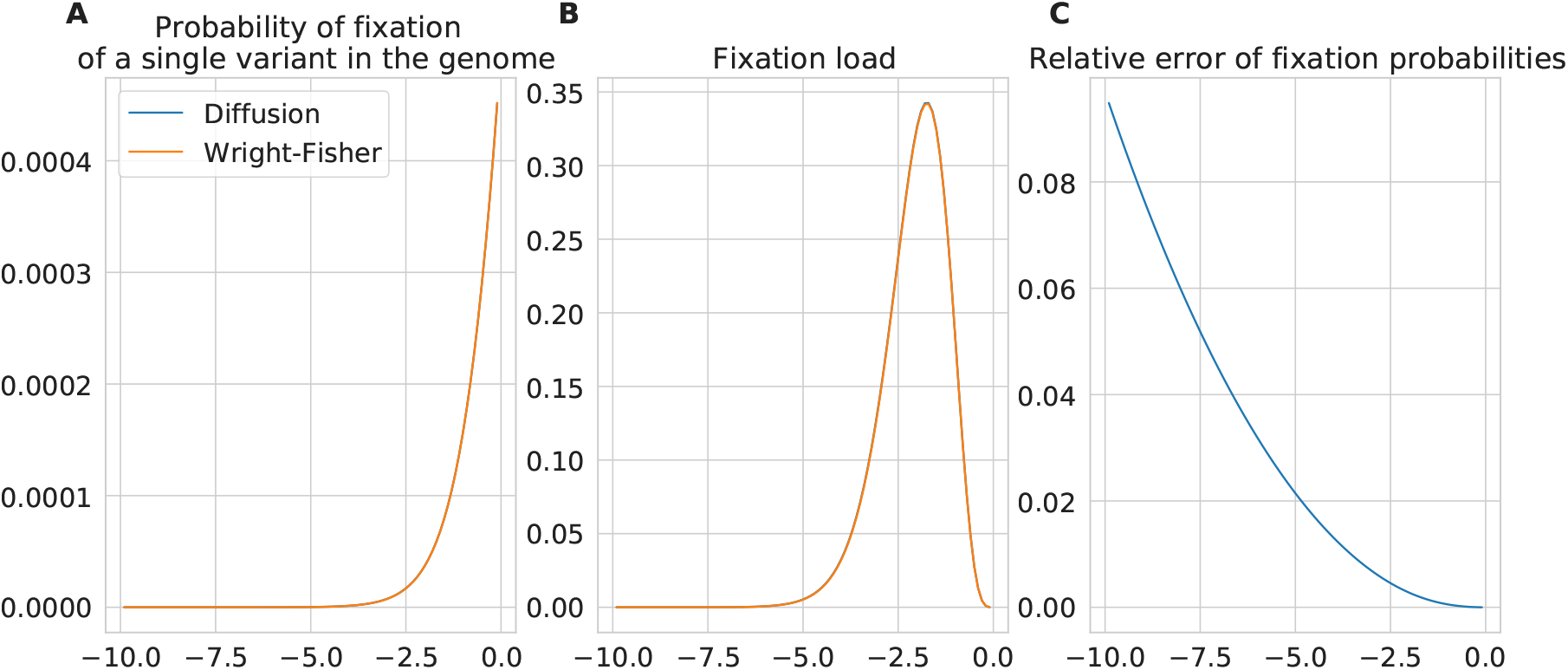
(A) Fixation probability, (B) fixation load, and (C) relative error between the diffusion and Wright-Fisher models for N=1000.

**Figure S5:**
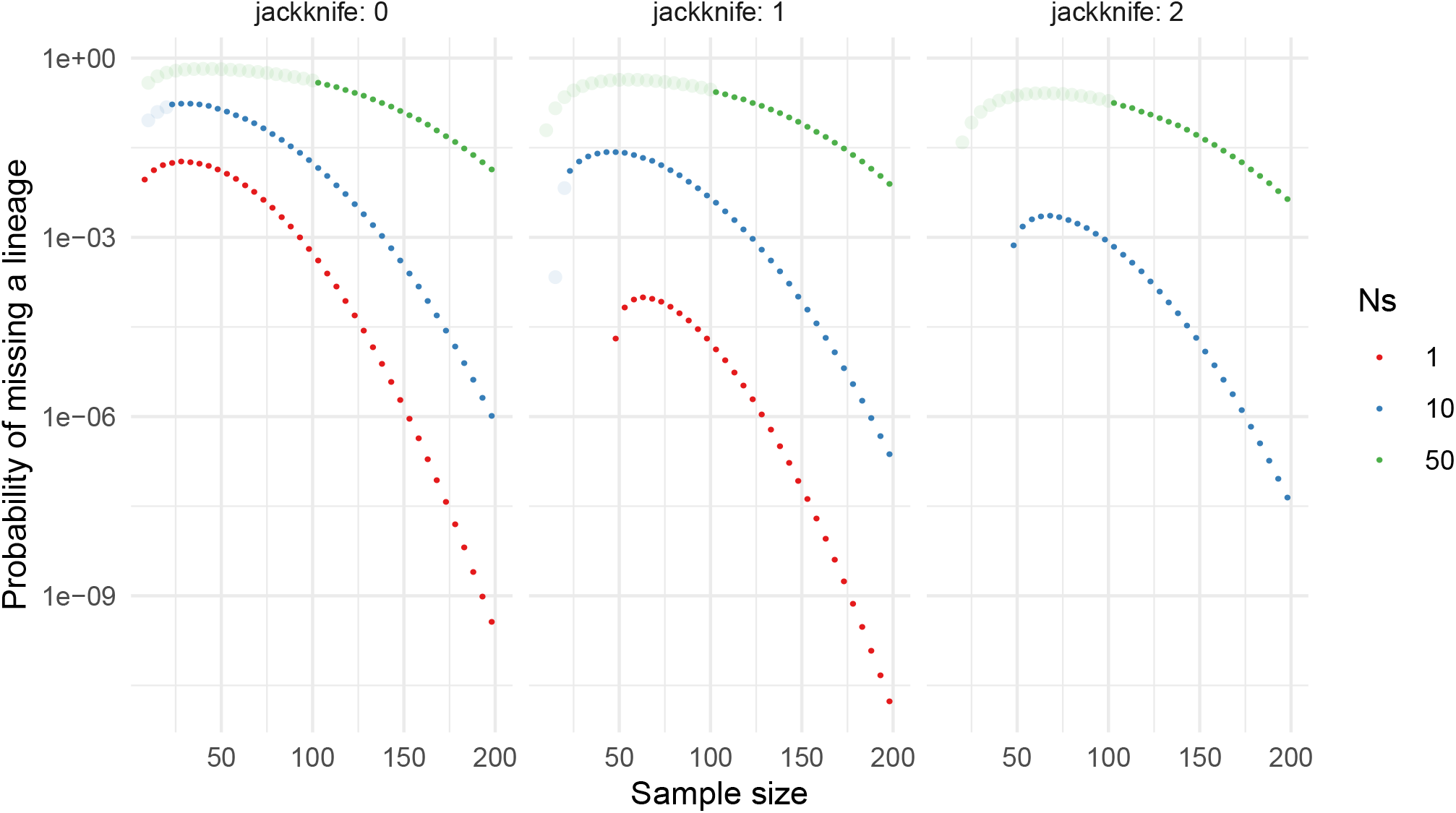
Probability that there are more contributing parents than offspring: *n_p_* > *n_o_* when every allele is derived (worst case). Calculated as 1 — Σ*_i_o__* **Q***_n_o__* (*i_p_,i_o_*), with *Q* defined in equation 4, with *N* = 1000. Each panel shows an increasing order of the jackknife approximation. Transparent points show sample sizes below the critical point, 2*Ns*.

**Figure S6:**
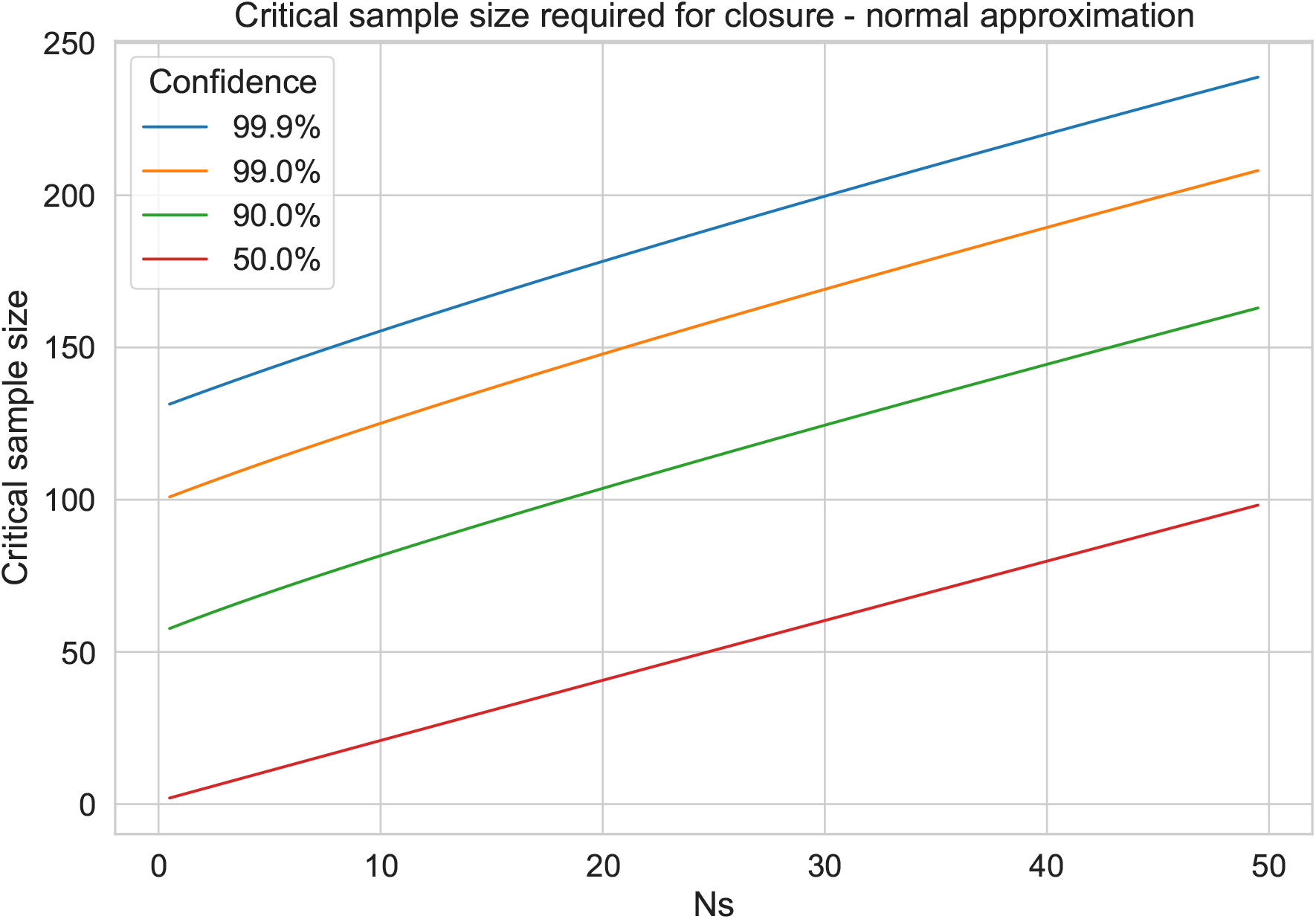
Critical sample size that is required to ensure that a given proportion of simulated draws are included, for a given closure confidence level. The 50^th^ percentile corresponds to the mean in the equation (8). Note that the normal approximation is invalid for *Ns* = 0, where the occupancy distribution should be used. All lines are calculated with *N_e_* = 1000.

**Figure S7:**
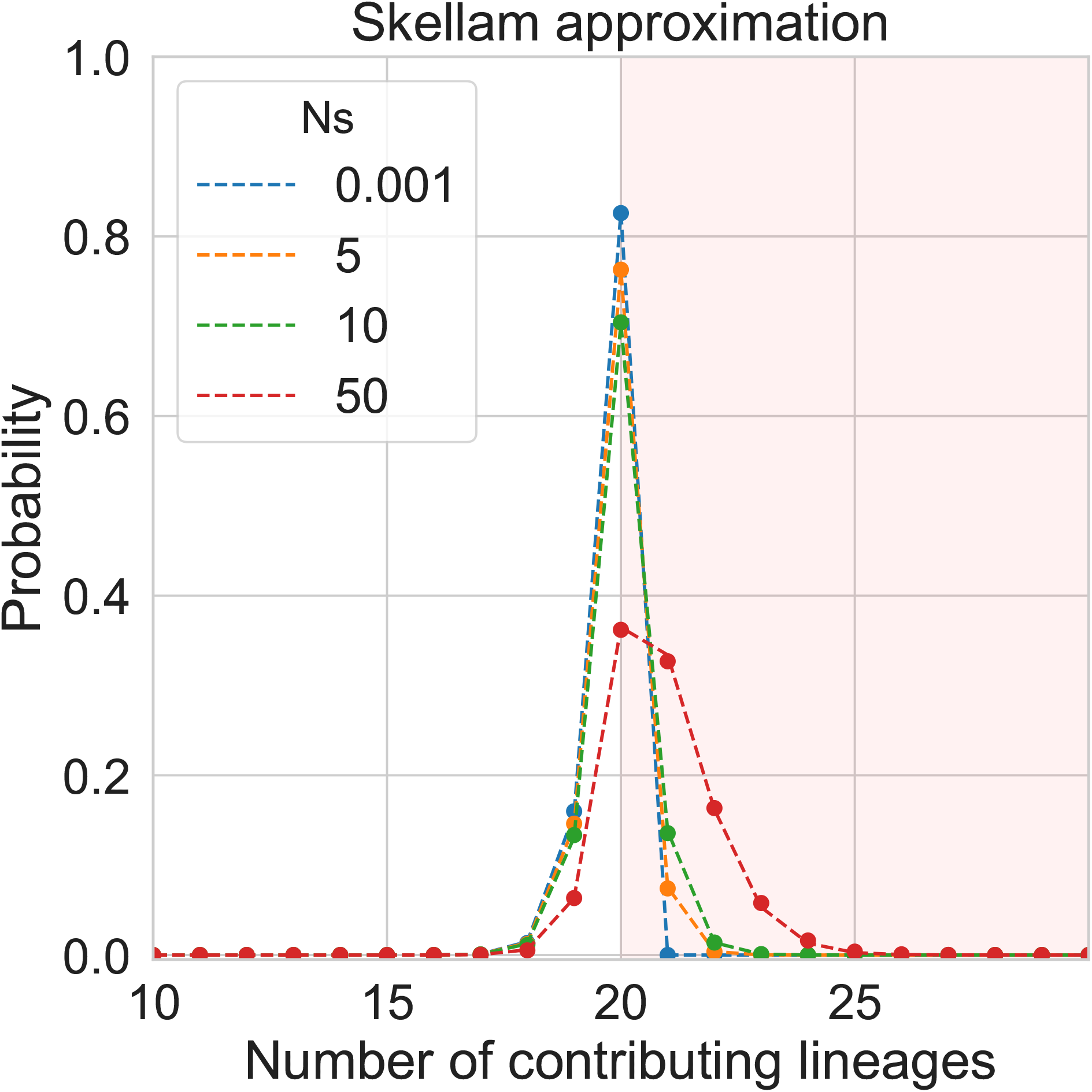
Skellam approximation to the number of required lineages where *n_o_* is small: 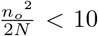. The Skellam distribution describes the difference between the number of offspring lineages *n_o_*, and parental lineages *n_p_*.

**Figure S8:**
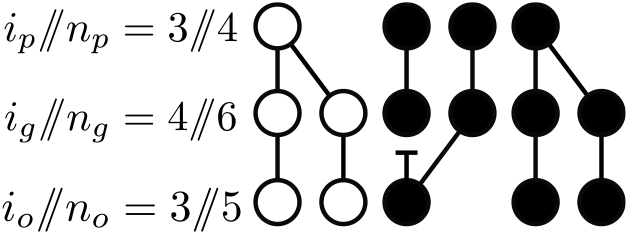
An extension of figure 1, with the addition of the gamete “generation”, with *n_g_* gametes, of which *i_g_* are derived.

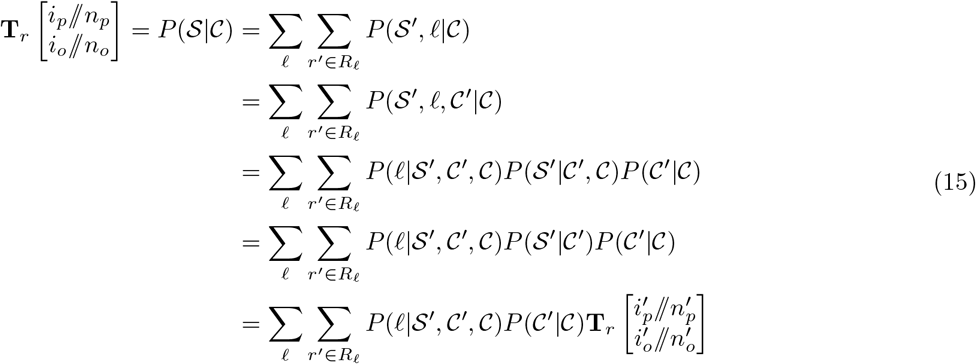

where the third line is an application of the chain rule and the fourth line uses the independence of primed events on the last draw. We now simply need to enumerate all the distinct types of events for the last draw and compute the corresponding probabilities, which we do on Figure 2.

### S1.3 Distribution on the number of parental lineages

#### S1.4 Distribution of number of sampled lineages

To disentangle the effects of drift and selection, in addition to tracking the parental and offspring lineages, we also define *n_g_* as the number of gametes sampled in the process (Fig. S8). The difference *n_g_ — n_o_* ≥ 0 depends on the number of selective deaths, whereas the difference *n_p_* — *n_g_* ≤ 0 depends on the number of coalescent events.

To get an analytical expression for the probability of missing lineages, in the worst-case scenario where all the parental alleles are derived, we consider the probability that *n_p_* parents have been sampled,

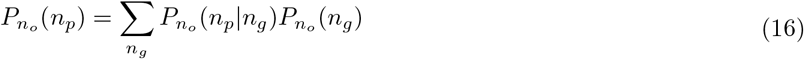

where *n_p_* and *n_g_* is the number of sampled parents and gametes, respectively (Fig. 1C).

The distribution over the number of gametes, *n_g_*, is given by the negative binomial, parameterized by the number of successes *n_o_*, and the probability of a successful draw is 1 — *s*.

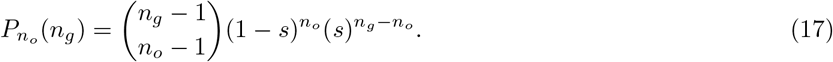

Given *n_g_*, the number of parental lineages *n_p_* follows the modified occupancy distribution (also known as the Arfwedson distribution) [Wakeley, 2009, O’Neill, 2019, Johnson et al., 2005]:

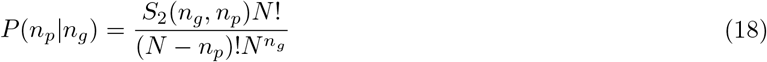

where *S*_2_(*n_g_, n_p_*) is a Stirling number of the second kind, which is the number of ways to partition *n_g_* gametes into *n_p_* parents (see Johnson et al. [2005] section 10.4 for a thorough treatment). The occupancy distribution requires exchangeability of the alleles, which is satisfied by the condition that all parental alleles are derived.

Combining the two distributions together through equation 16, we get:

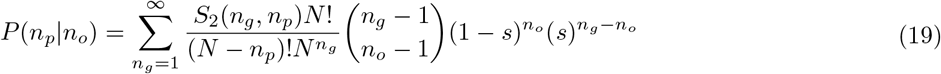

**Figure S9:**
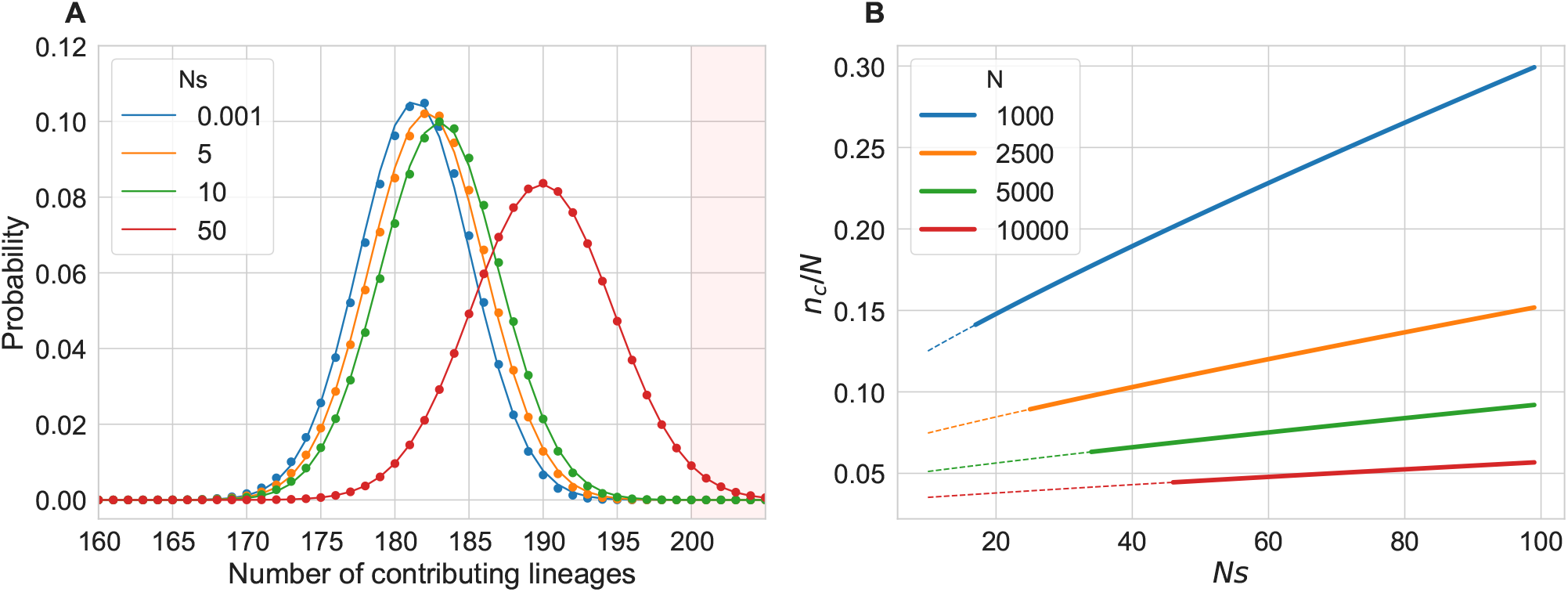
**A** The distribution of number of required lineages with *n*_0_ = 200. Shaded red area shows missing probability, where *n_p_ > n_o_*. Points represent the exact probability one generation into the past (Eq. 19), solid lines - Gaussian approximation. **B** Sample size (*n_o_*) as a fraction of population size (*N*) such that we have 99% confidence that no lineages are missing, derived from the Gaussian approximation. The dotted line indicates regimes where the Gaussian approximation is likely inaccurate. In both panels, *N* = 1000.

We did not find an analytical expression for this sum, but it can be computed efficiently using methods presented in [O’Neill, 2019]. Figure S9A (dotted line) shows the distribution of the number of contributing parental lineages for several selection coefficients with *n_o_* = 200. As the strength of selection is increased, we begin requiring larger numbers of lineages, while with increasing sample size, there is a decrease in required lineages due to coalescent events.

### S1.5 Gaussian approximation

The distribution in equation 19 can quickly be calculated numerically, but provides little intuition. We therefore compute the expectation and variance of this distribution, and show that it can be accurately approximated by Skellam and Gaussian distributions in useful parameter regimes.

Using the law of total expectation, we can write the expectation *E*[*n_p_* — *n_o_*|*n_o_*] as

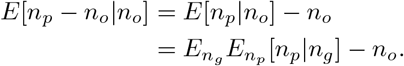

The expectation over *n_p_* is simply that of the occupancy distribution Wakeley [2009].

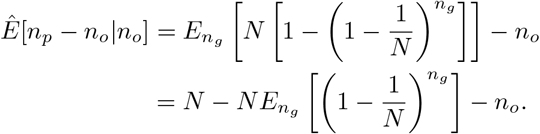

As mentioned above, the number of selective deaths, *n_g_ — n_o_* follows a negative binomial distribution with success probability 1 — *s*. We can use the moment generating function of the negative binomial to compute the expectation of *k^n_g_^* for any constant *k*:

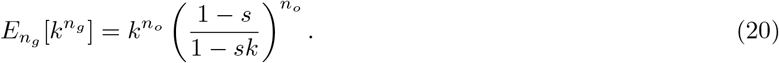

Thus we can write:

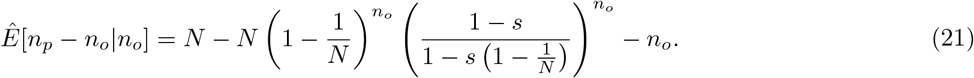

Taking the terms of order up to 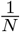 gives

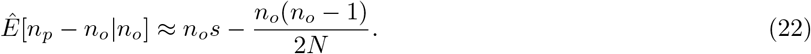

We thus have the usual interplay of selection and drift. Solving for 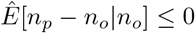 yields, to leading order,

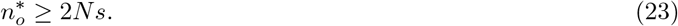

This represents a critical sample size (Fig. S6) where there is more drift than selection events, on average. The next section outlines a similar approach to compute the variance of this distribution, Var[*n_p_ — n_o_*|*n_o_*].

#### S1.5.1 Variance of number of contributing lineages

We can obtain the variance of the distribution of the number of parental lineages *P*(*n_p_*|*n_o_*) by using the law of total variance:

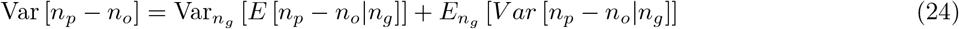

As previously, we are assuming that all the parental alleles are derived. The expectation in the first term can be derived from the occupancy distribution and the identity 20:

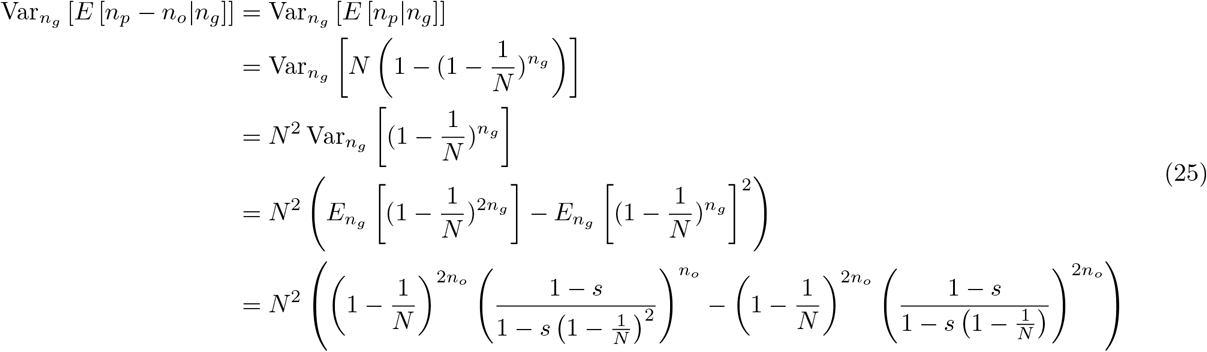

The variance of the second term of Equation (24) is the variance of the modified occupancy distribution Johnson et al. [2005]:

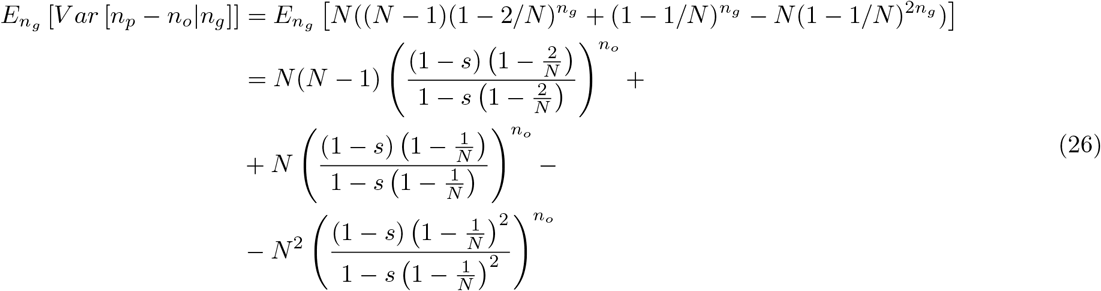

Combining the two terms in Equation (24), we get

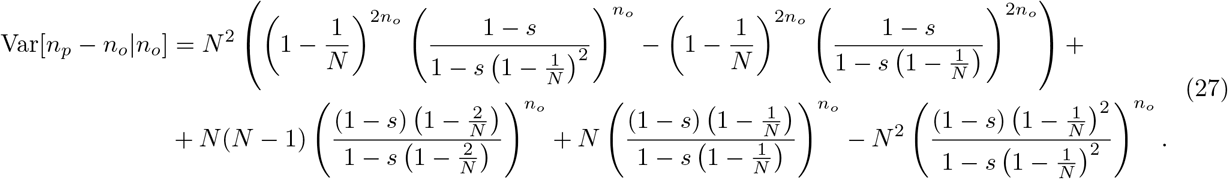

Taking series expansion in Mathematica [Wolfram Research, Inc., 2020], we can show that

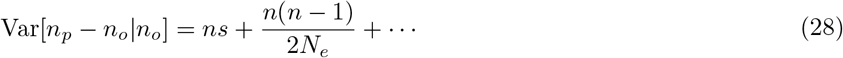

where missing terms are of at least second order in products of 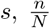, and 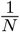. The code is available in the github repository.

### S1.6 Simplifying Gaussian approximation

Given a *z* for the change in the number of lineages of 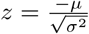 we use 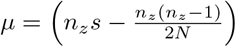 and 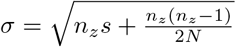 from Equations 7 and 9 to obtain a bound for *n_z_*:

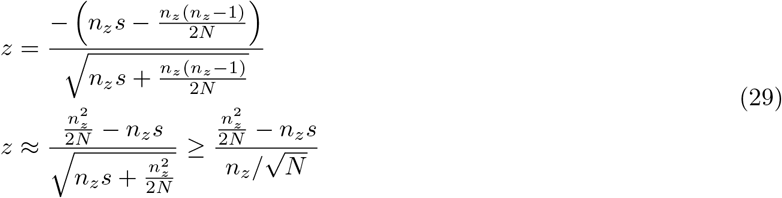

The latter inequality uses the fact that we are in the regime where the expected number of drift events is larger than the number of selection events. Thus 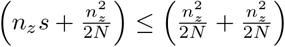.

This can now be trivially solved for *n_z_* to yield

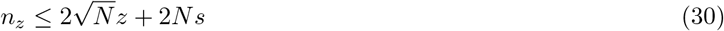

### S1.7 Construction of transition probability matrices with multiple generations

Multi-generation transition matrices can simply be computed by iterating equation (3). Assuming, for simplicity, that the population sizes and selection coefficients are constant over the last two generations, and using matrix products to account for sums over the number *i_p_* of parental derived alleles, we find

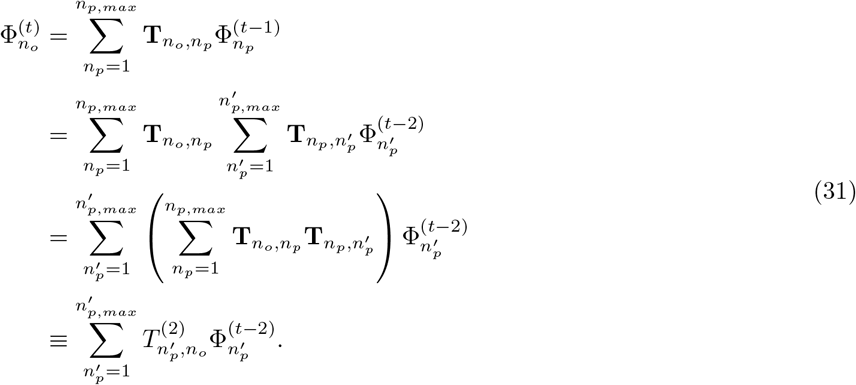

where 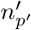, is the number of sampled lineages in the grandparental generation, and *n_p,max_* and 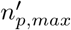 are the maximum number of sampled parental alleles considered in the parental and grandparental generations, respectively. We can choose *n_p,max_* > *n_o_* and *n_p,max_* = *n_o_* to preserve closure of the moment equation while allowing for drift to cancel out selection events over the course of two generations.

Since each matrix product takes *O*(*n_o_n_p_n_p’_*), and there are at most *O*(*n_p,max_*) such products, the computation of this matrix product takes at most 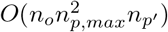. This scaling can be improved upon in numerical implementations by summing only over values of *n_p_* that contribute appreciably. In addition, we need to build the matrices 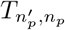 themselves. In the recursive approach, the computation of the matrix for the largest sample includes the computation of matrices for smaller sample sizes, so the computation time is at most that of the largest possible matrix, 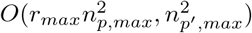. In the exact formulation of our model, *n_p_* can be as large as *r_max_n_o_*, however this would require every draw to be rejected by selection and only contribute terms of order *s^n_o_r_max_^*. A numerically appropriate cutoff for a given *n_o_* and *s* can be computed dynamically by keeping track of the proportion of unaccounted-for lineages. In most practical applications with *s* < 0.1 we expect that choosing, e.g., *n_p,max_* = 2*n_o_* would provide excellent convergence, hence an overall scaling of 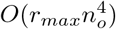 for the construction of the matrices and 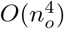 for the matrix product. Thus a naive construction of a transition matrix over *g* generations would require 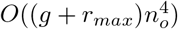.

